# Interrupting reactivation of immunological memory reprograms allergy and averts anaphylaxis

**DOI:** 10.1101/2020.03.04.975706

**Authors:** Kelly Bruton, Paul Spill, Shabana Vohra, Owen Baribeau, Saba Manzoor, Siyon Gadkar, Malcolm Davidson, Tina D. Walker, Yosef Ellenbogen, Alexandra Florescu, Jianping Wen, Derek K. Chu, Susan Waserman, Rodrigo Jiménez-Saiz, Slava Epelman, Clinton Robbins, Manel Jordana

## Abstract

IgE production against innocuous antigens can lead to life-threatening reactions such as anaphylaxis. While IgE levels drastically decline with strict allergen avoidance, the ability to regenerate IgE can persist for a lifetime, as is the case for peanut allergy. The mechanism by which IgE regenerates remains unresolved. A novel culture system and application of single-cell RNA-sequencing, elucidated the transcriptomic signature of human peanut-reactive B and T cells and revealed IL-4/IL-13 as a signaling pathway critical for the IgE recall response. Indeed, interruption of this pathway not only prevented IgE production and anaphylaxis, but also reprogrammed the pathogenic response against peanut. This investigation advances our understanding of the mechanism that regenerates IgE in food allergy and spotlights IL-4/IL-13 blockade as a therapeutic with disease-transforming potential.

**One Sentence Summary:** Single-cell transcriptomics of allergic memory responses reveals how to teach the immune system to forget.

## Main Text

Adaptive immunity’s hallmarks of specificity and memory play a central role in human health and disease. Studying pathogen-host interactions have revealed fundamental immunological insights and inform the rational design of therapeutics (*e.g.* vaccines). However, maladaptive immune responses, such as in allergy, may not operate under the same rules as in infection. Therefore, understanding the mechanisms of allergic disease may uncover novel immunological principles and also inform new potential therapies.

Food allergy affects millions worldwide and remains devoid of disease-transforming therapies. Immunoglobulin (Ig) E is the key effector molecule mediating food-induced allergic reactions. While there is evidence that serum levels of allergen-specific IgE drastically decline after extended periods of allergen avoidance (*1*–*3*), the capacity to regenerate IgE following re-exposure to allergens such as peanut (PN), tree nuts and shellfish remains for a lifetime, thus posing a perpetual risk of life-threatening systemic reactions (anaphylaxis). It is now thought that the lifelong capacity to regenerate IgE resides in the persistence of allergen-specific memory B cells (MBCs). The extreme rarity of IgE^+^ MBCs (*4, 5*) and the predisposition for IgE to be derived through sequential class-switching from an IgG intermediate (*6*) have positioned IgG^+^ MBCs as the chief reservoir for IgE regeneration (*3, 7*). Here, we investigated PN allergy in humans and mice using a novel model system, comprehensive protein analysis, and single-cell transcriptomics to better understand mechanisms of Th2 cellular and humoral immune memory. Our findings illustrate that interrupting reactivation of immunological memory aborts the re-emergence of IgE and fully prevents anaphylaxis through sustained reprogramming of pathogenic Th2 responses.

The severity of symptoms to PN exposure in allergic individuals prevents the study of underlying mechanisms that regulate recall of harmful immune responses in a clinical setting. To circumvent this issue, we developed an *in vitro* platform utilizing human peripheral blood mononuclear cells (PBMCs) from PN-allergic patients to interrogate allergen-dependent T and B lymphocyte responses (Fig. 1A and Table S1). A targeted proteomic analysis of supernatant from PN-stimulated PBMCs in 7 PN-allergic donors revealed upregulation of canonical Th2 cytokines, including IL-5, IL-9, and IL-13 (Fig. 1B and C and Fig. S1). *In vitro* treatment of PBMCs with recombinant IL-4 and anti-CD40 resulted in IgE production detected at the cellular, molecular, and transcriptional level (Fig. S2A-C), irrespective of allergic status of donor cells. However, IgE responses to PN occurred in PBMCs from PN-allergic patients only; *in vitro* exposure of cells to PN increased numbers of IgE secreting cells (Fig. 1D), elevated levels of PN-specific IgE in culture supernatants (Fig. 1E), and increased transcription of *IGHE* (Fig. 1F). The extent of IgE produced *in vitro* directly correlated with PN-specific IgE levels detected in serum of PN-allergic donors (Fig. S2D-F).

**Fig. 1.**
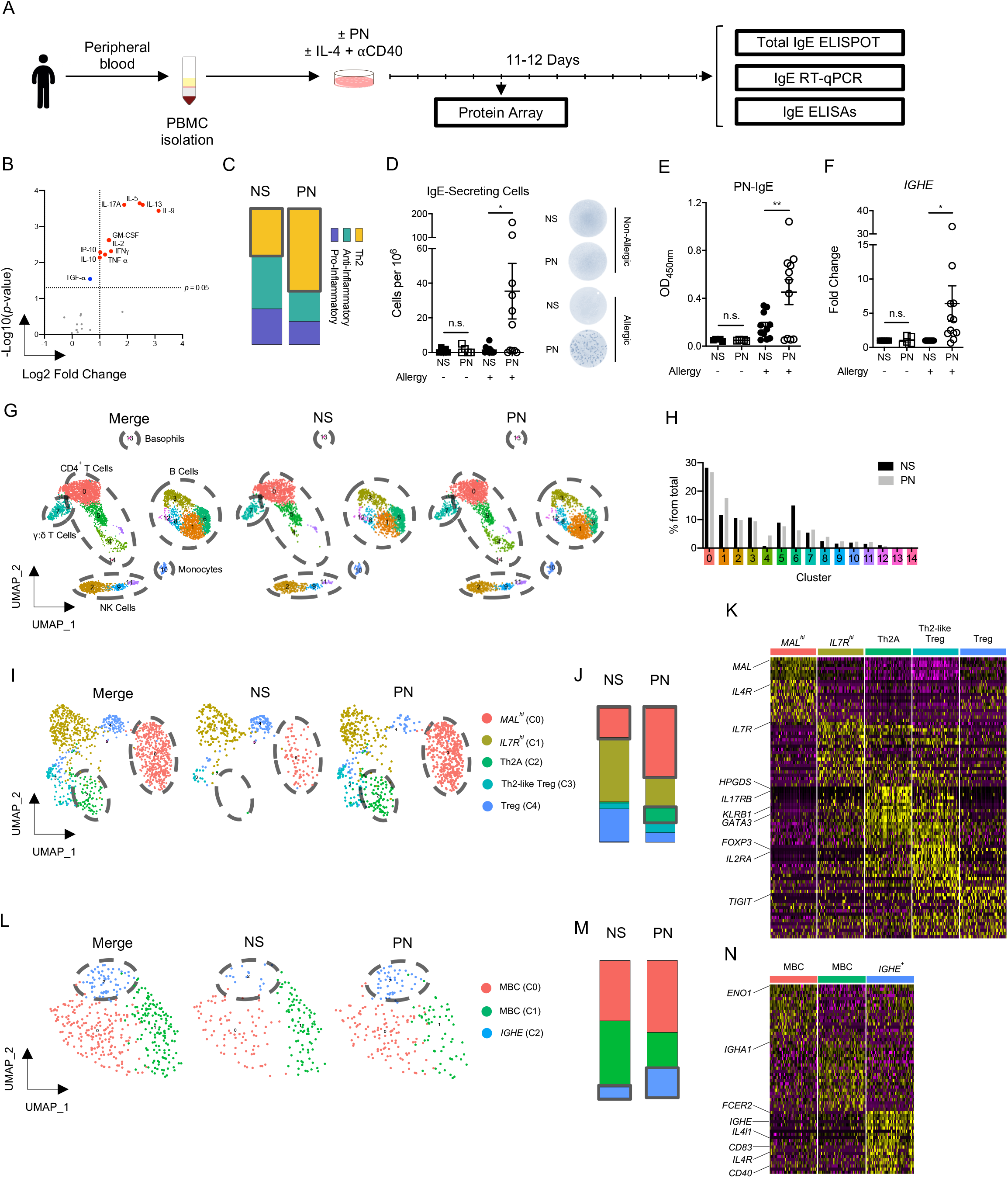
Characterization of recall response to PN in human PBMCs. (A) Schematic of culture with PN-allergic or non-allergic donor PBMCs. (B) Volcano plot of cytokines detected with 38-analyte array in PN-stimulated PBMCs from PN-allergic donors. Benjamini-Hochberg (BH) correction was applied to correct for multiple comparisons. Data are plotted as upregulated cytokines (red), downregulated cytokines (blue), and unchanged (grey). (C) Stacked bars comparing cytokine production, based on putative functional class. (D) Enumerated IgE-secreting cells detected at end of culture with representative images. (E) PN-specific IgE measured from culture supernatant. (F) Fold change of *IGHE* mRNA expression relative to housekeeping gene (*18S*) and non-stimulated control. Data are representative of 5 non-allergic donors (D-F) and 7-11 PN-allergic donors (B-F) and are plotted as mean ± SEM (D-F). \**P* < 0.05, \*\**P*<0.01. (G) UMAP identifying distinct leukocyte populations from a PN-allergic donor. (H) Frequency of cells in each cluster. (I) UMAP of subclustered T cells. (J) Stacked bars of subcluster frequency in I. (K) Heatmap of top 20 differentially expressed genes in T cell subclusters. (L) UMAP of subclustered MBCs. (M) Stacked bars of subcluster frequency in L. (N) Heatmap of top 20 differentially expressed genes in MBC subclusters.

To assess the transcriptional response of CD4 T and B cells to PN, we performed single-cell RNA-sequencing (scRNA-seq) on PBMCs isolated from non-stimulated (NS; 3272 cells) and PN-stimulated (3787 cells) culture from a representative PN-allergic donor (Fig. S3A). After quality control, removal of low-quality cells and doublets, Uniform Manifold Approximation and Projection (UMAP) was used to reduce the number of dimensions and visualize cell clustering. Assigning nomenclature derived from previously annotated gene signatures revealed 15 discrete cell clusters (C), including populations of CD4 T cells, B cells, NK cells, monocytes, and basophils (Fig. 1G and H and Fig. S3B) with (Fig. S3C). Sub-clustering of T cells expressing *GATA3, FOXP3, IL7R*, and *MAL* (Fig. S4A) identified 5 cell populations (Fig. 1I and Table S2). C0 expanded considerably upon PN stimulation (Fig. 1J), expressed high levels of *MAL* (Fig. 1K), and presumably responds to IL-4 signaling during early commitment to the Th2 lineage (*8*). Accordingly, C0 cells uniquely expressed *IL4R* (Fig. S4B) and were enriched for genes with annotated functions related to IL-4 signaling and activation (Fig. S4C). However, the presence of C0 T cells even under baseline (NS) conditions questions their specificity and functional relevance. C2 was uniquely identified in PN-stimulated PBMCs (0.8% NS to 10.6% PN) (Fig. 1I and J). Analysis of differentially expressed genes (Table S2) defined an activated Th2 phenotype (*GATA3, IL17RB, KLRB1, IL2RA*, and *CTLA4*), high expression of prostaglandin D synthase (*HPGDS*), and the prostaglandin D2 receptor, CRTH2 (*PTGDR2*) (Fig. 1K). The data identify a transcriptional profile consistent with a recently identified population of Th2 cells that drive clinical allergic disease, coined as “Th2A” cells (*9*).

Subclustering MBCs, which we identified by expression of the memory markers *CD27* and *AIM2* and low expression of *IGHM, IGHD*, and *TCL1A* (*10*) (Fig. S5A and B and Fig. 1L), identified 3 distinct clusters. C0 and C1 contained heterogeneous IgH expression but did not express *IGHG4* and *IGHE* (highly expressed in C2 cells) (Fig. S5C). *IGHE*-expressing C2 cells expanded with PN stimulation (Fig. 1M). *IGHE*^*+*^ cells were transcriptionally unique (Fig. 1N and Table S3), expressing genes associated with co-stimulation (*e.g. CD40* and *CD83*) and IL-4-signaling (*11, 12*) *(e.g. IL4l1* and *GCSAM)*, and high expression of *IL4R* (Fig. S5D). In sum, we have developed an *in vitro* platform that yields robust Th2 humoral responses with PN stimulation, positioning this as a novel system by which to study the human allergic recall response. Application of scRNA-seq to *in vitro* stimulated PN-allergic PBMCs revealed the transcriptomic profile of PN-reactive B and T cell subsets. Cell populations that expanded with PN stimulation exhibited an upregulation of genes related to IL-4 signaling.

Given the enrichment of IL-4 signaling profiles among PN-reactive B and T cell clusters, we next assessed the mechanism of human memory IgE production by blocking IL-4Rα during recall. IL-4Rα is a shared chain between IL-4 and IL-13 receptors and thus, its blockade prevents IL-4 and IL-13-signaling. Anti-IL-4Rα remarkably inhibited IgE responses, including a reduction of IgE-secreting cells (Fig. 2A), decrease of PN-specific IgE production (Fig. 2B), and prevention of *IGHE* transcription (Fig. 2C). Suppression of IgE responses did not associate with apoptosis of CD4 T cells or B cells through deprivation of IL-4 signaling (Fig. S6). Instead, anti-IL-4Rα treatment downregulated typical Th2-associated cytokines including, IL-5 and IL-9, but increased production of IL-10, IL-1α, IL-1β, MIP-1α, G-CSF, and IFNxsγ (Fig. 2D-E and Fig. S7). The skewing towards a Th1 and IL-10 dominant phenotype (Fig. 2F) suggests cellular “reprogramming” of memory.

**Fig. 2.**
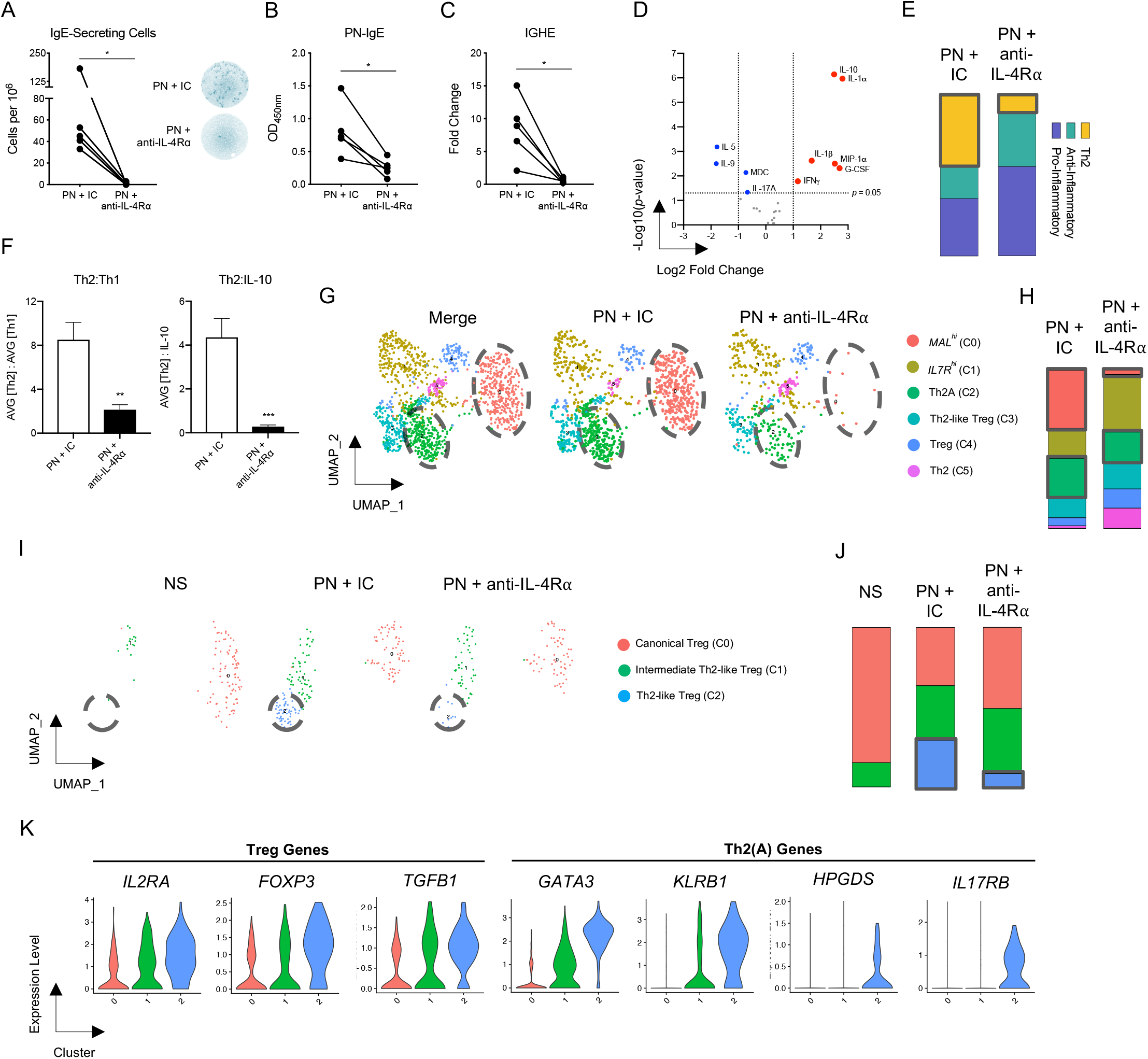
IL-4/IL-13 signaling blockade aborts IgE recall response and skews Th2 polarization in human PBMCs. (A) Enumerated IgE-secreting cells detected at end of culture with representative images. (B) PN-specific IgE measured from culture supernatant. (C) Fold change of *IGHE* mRNA expression relative to housekeeping gene (*18S*) and non-stimulated control. (D) Volcano plot of cytokines detected with 38-analyte array in IC or anti-IL-4Rα-treated PN-stimulated PBMCs from PN-allergic donors. Benjamini-Hochberg (BH) correction was applied to correct for multiple comparisons. Data are plotted as upregulated cytokines (red), downregulated cytokines (blue), and unchanged (grey). (E) Stacked bars comparing cytokine production, based on putative functional class. (F) Ratio of Th2 cytokines to Th1 or IL-10. Data are plotted as mean ± SEM (C) or individual donors (D-F). \**P* < 0.05, \*\**P*<0.01, \*\*\**P*<0.001. (G) UMAP of subclustered T cells. (H) Stacked bars of subcluster frequency in G. (I) UMAP of subclustered Tregs. (J) Stacked bars of subcluster frequency in I. (K) Violin plots depicting gene signature in Treg subclusters.

To better understand this, we used scRNA-seq to compare the profiles of over 6000 human PN-stimulated PBMCs with or without anti-IL-4Rα treatment (3427 PN + isotype control (IC) and 2718 PN + anti-IL-4Rα). The PN-induced expansion of the *MAL*^*hi*^ population was completely inhibited by IL-4Rα blockade (Fig. 2G and H). This was consistent with their high expression of *IL4R* and suggest that IL-4/IL-13 signaling is critical for their expansion. In contrast, IL-4/IL-13 signaling was dispensable for maintaining terminally differentiated Th2A cells (C2). Sub-clustering of Tregs, which have previously been shown to acquire a Th2-skewed phenotype in allergic disease (*13*), revealed 3 clusters (Fig. 2I and J): canonical Tregs (C0), the dominant population in NS PBMCs, and 2 clusters of Th2-skewed Tregs (C1 and C2) differing in intensity of Th2 gene expression, including *GATA3, HPGDS*, and *IL17RB* (Fig. 2K). Anti-IL-4Rα specifically attenuated PN-induced emergence of C2 Th2-like Tregs, suggesting a role of IL-4/IL-13 signaling in Treg reprogramming. With regards to MBCs, the expansion of *IGHE*^+^ cells was abrogated by IL-4Rα blockade, which coincides with upregulation of IL-4-signaling genes in this cluster (Fig. S8). These data establish the critical requirement of IL-4/IL-13 signaling in human IgE memory responses to PN and the capacity of IL-4Rα blockade to induce non-Th2 skewing in CD4 T cell subsets, including Tregs.

To investigate the impact of IL-4Rα blockade *in vivo*, we employed a murine model of epicutaneous allergic sensitization (*14*). This produced allergen-specific IgG1 and IgG1^+^ MBCs, with IgE produced only upon non-sensitizing, subcutaneous (*s.c.*) allergen re-exposure (Fig. 3A). We took advantage of the temporal control of IgE production and treated mice with anti-IL-4Rα immediately before *s.c.* re-exposures. Emergence of IgE was fully abrogated by anti-IL-4Rα treatment (Fig. 3B). Upon challenge, mice receiving anti-IL-4Rα exhibited decreased clinical reactivity, including reduced drop in core body temperature (Fig. 3C), lower hematocrit levels (Fig. 3D), and reduced severity of clinical symptoms (Fig. 3E). We reasoned that this partial response could be a consequence of IgG-mediated alternative anaphylactic pathways, which have been observed to operate more so in mice than in humans. To this end, mice were treated with anti-CD16/32 before challenge to block IgG binding to Fcγ receptors (*15*). Mice co-treated with anti-IL-4Rα and anti-CD16/32 displayed no clinical reactivity upon challenge, as evidenced by a maintenance of homeostatic core body temperature (Fig. 3C) and hemoconcentration (Fig. 3D).

**Fig. 3.**
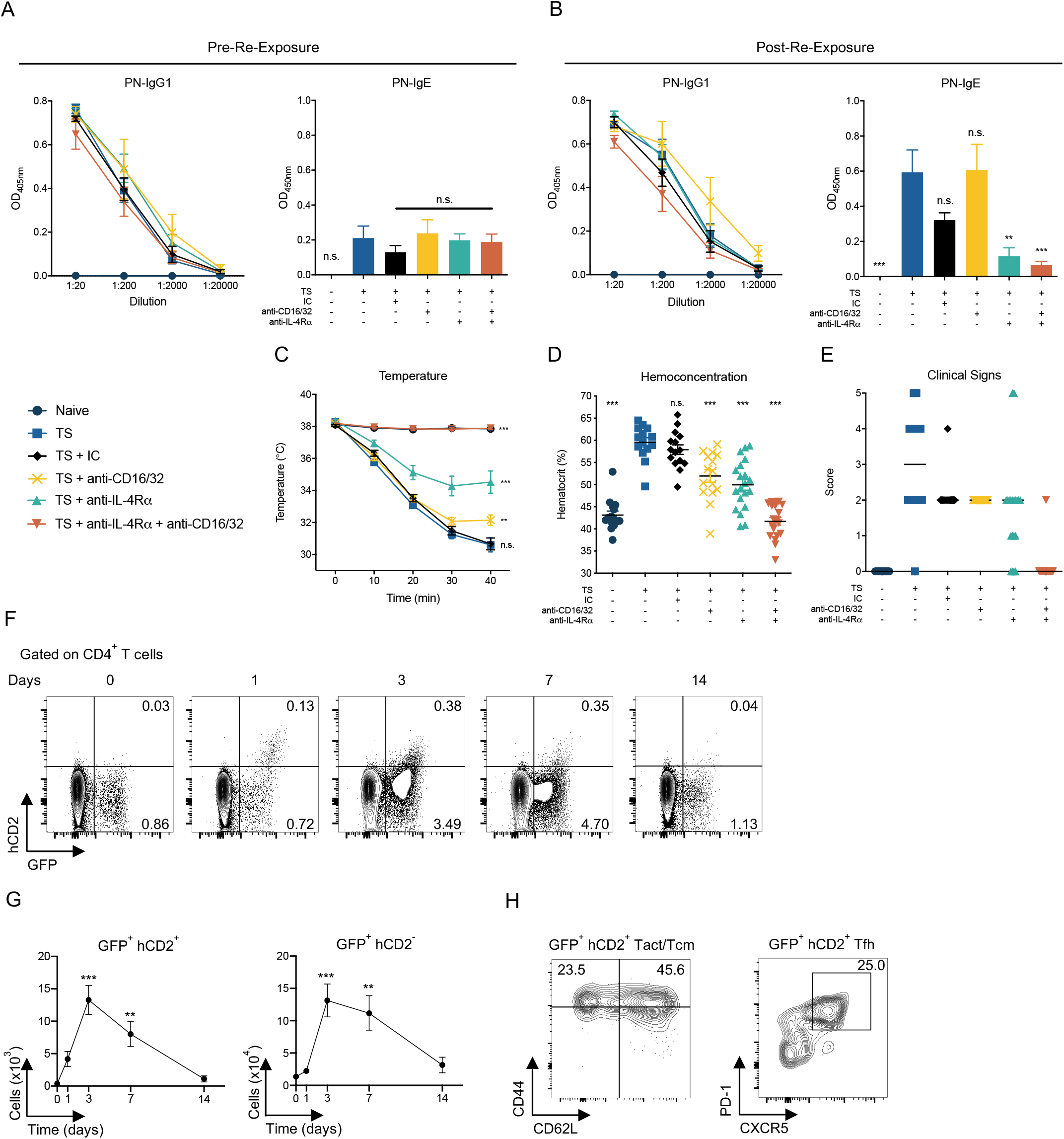
*In vivo* model reveals requirement of IL-4/IL-13 signaling and source of IL-4 in recall response. (A) Serum PN-specific IgG1 and IgE following sensitization. (B) Serum PN-specific IgG1 and IgE following re-exposure. (C) Core body temperature at 10-minute intervals during systemic challenge. (D) Hemoconcentration at 40-minutes post-challenge. (E) Scoring of clinical signs during systemic challenge. (F) Representative contour plots of *IL4*-transcribing and IL-4 secreting CD4^+^ T cells 0-14 days following PN re-exposure with averaged frequencies (in right quadrants). (G) Absolute counts of *IL4-*transcribing and IL-4-secreting CD4^+^ T cells. (H) Representative contour plot and averaged frequency of CD44 + CD62L and PD-1 + CXCR5 expression in IL-4-secreting CD4^+^ T cells at 3 days post-re-exposure. Data are representative of 2-3 experiments with 3-5 mice per experimental group, plotted as mean ± SEM or median. \*\**P*<0.01, \*\*\**P*<0.001.

To elucidate the cellular source of IL-4 that drives the memory recall response *in vivo*, we sensitized IL-4 dual-reporter (4get/KN2) mice to PN, as before. The approach enabled discrimination between cells expressing *IL4* transcripts (GFP^+^ hCD2^-^) and cells actively secreting IL-4 (GFP^+^ hCD2^+^) (*16*). Despite a haploinsufficiency for *IL4* (*17*), sensitized 4get/KN2 mice developed PN-IgE upon re-exposure and were clinically reactive, albeit at lesser degree than wild-type counterparts (Fig. S9A-C). We found that at all time points up to, and inclusive of, 7 days post-re-exposure, CD4 T cells averaged over 92% of IL-4-secreting cells in the draining lymph nodes (Fig. S9D and E). GFP^+^ hCD2^+^ and GFP^+^ hCD2^-^ cells peaked at day 3 post-re-exposure with contraction by day 14 (Fig. 3F and G). Among IL-4-secreting CD4^+^ T cells, 46% co-expressed CD44 and CD62L, suggesting a central memory (Tcm) phenotype and 25% exhibited a canonical T follicular helper (Tfh) cell phenotype (Fig. 3H). Thus, antigen-experienced CD4^+^ T cells provide the overwhelming source of IL-4 in a recall response, with little to no contribution from other cell types. In sum, IL-4Rα blockade aborts IgE production and prevents anaphylaxis, suggesting that reactivation of memory is dependent on CD4 T cell-derived IL-4.

Lastly, we investigated whether anti-IL-4Rα-mediated effects were only operative in presence of the antibody (*i.e.* transient) or if sustained immunological reprogramming was induced. To this end, mice were epicutaneously sensitized, treated with anti-IL-4Rα, and re-exposed as earlier described, then we allowed 6 weeks for anti-IL-4Rα clearance before re-exposing mice 3 additional times to PN (Fig. 4A). Consistent with previous experiments, IgG1 was detectable following sensitization (Fig. 4B) and IgE production was induced after *s.c.* re-exposure in untreated allergic mice, but not in anti-IL-4Rα-treated allergic mice (Fig. 4C). Following discontinuation of anti-IL-4Rα treatment, we observed a sustained and complete inhibition of IgE production despite antibody clearance (Fig. 4D). Therefore, suppression of IgE production by anti-IL-4Rα persists long-term, identifying a previously unrecognized plasticity of IgE humoral immunity with novel therapeutic potential to redirect the allergic response.

**Fig. 4.**
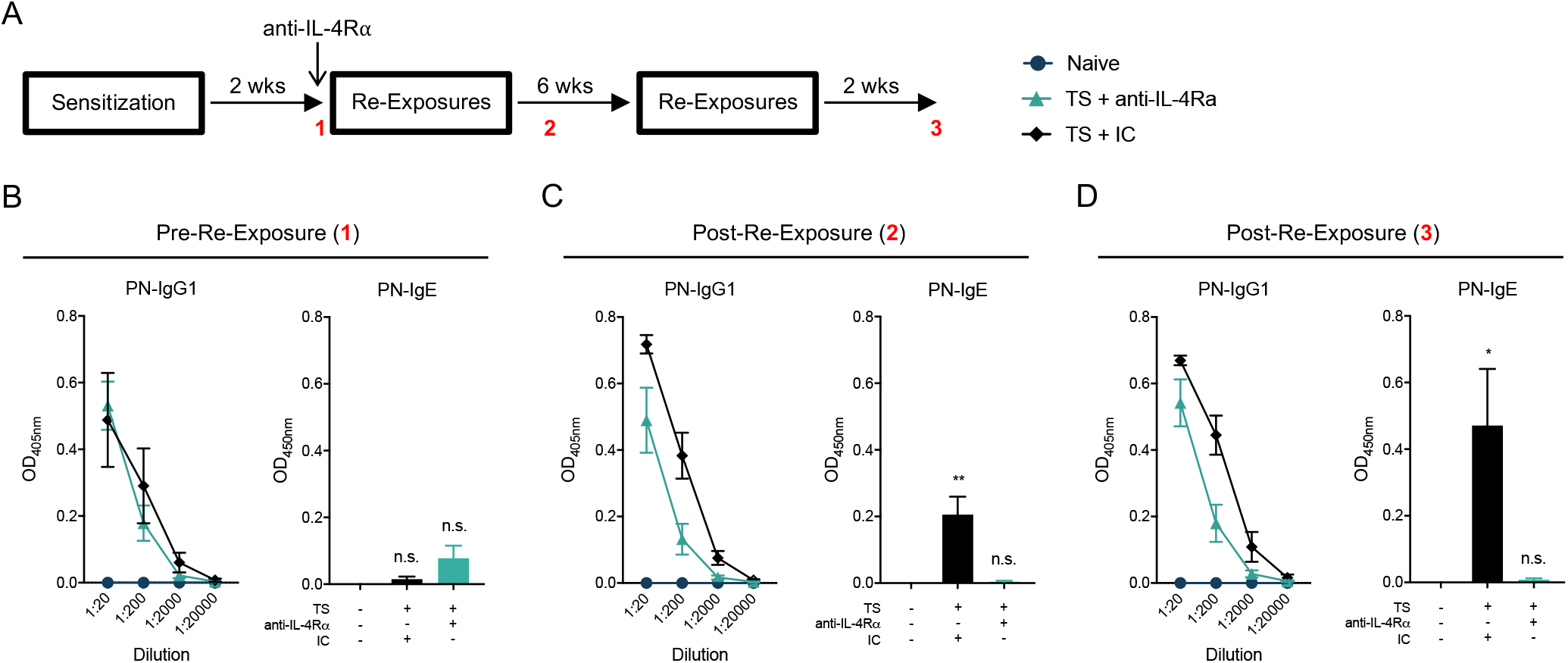
Inhibition of IgE production is sustained following anti-IL-4Rα clearance. (A) Experimental schematic. Red numbers depict timepoint of blood collection. Serum PN-specific IgG1 and IgE following sensitization (B), re-exposure with anti-IL-4Rα (C), and re-exposure without anti-IL-4Rα (D). Data are representative of 2 experiments with 5 mice per experimental group, plotted as mean ± SEM. \**P*<0.05, \*\**P*<0.01.

IgE effector immunity requires constant memory reactivation for persistence. This perspective distinguishes IgE responses from classical paradigms of humoral Th1 immune memory to pathogens, where a single exposure can elicit protection for decades. Combining protein profiling, single cell transcriptomics, and *in vitro* and *in vivo* assays, our findings usher a paradigm whereby Th2 immune memory is activated by CD4 T cell-derived IL-4, yielding the emergence of *MAL*^*hi*^ CD4 T cells, Th2A cells, and IgE-secreting cells. This allergic response can be reprogrammed by transient interruption in IL-4/IL-13 signaling at the time of memory reactivation. Altogether, the data have implications for conceptualizing therapeutic manipulation of IgE responses.

Clinical trials employing anti-IL-4Rα monoclonal antibodies have proven beneficial in reducing disease activity in atopic dermatitis, eosinophilic asthma, and chronic rhinosinusitis with nasal polyposis (*18, 19*). However, the mechanism of action remains largely unexplored, particularly in terms of downstream effects on memory. We provide direct evidence regarding the therapeutic potential of anti-IL-4Rα in the context of established PN allergy, where interruption of IL-4/IL-13 signaling inhibits IgE recall responses, skews the Th2-dominant cytokine response, prevents induction of anaphylaxis, and reprograms the immune response against PN.

## Materials and Methods

### Study Population

A cohort of 12 PN-allergic and 5 non-allergic blood donors were recruited from McMaster University and McMaster Children’s Hospital (Hamilton, ON). Allergy to PN was determined by PN-specific IgE ImmunoCap^®^ performed at LRC Hamilton (McMaster Children’s Hospital), by saline and histamine-controlled skin prick test, and clinical history of reactivity. PN-allergic individuals were considered for inclusion with PN-specific serum IgE levels >0.35 kU/L and skin prick test ≥3 mm greater than saline control. All allergy testing was performed no earlier than 6 months before blood was drawn for experimentation. Exclusion criteria for all recruited donors included: allergen immunotherapy, previous or current omalizumab (Xolair^®^), dupilumab (Dupixent^®^), or other systemic immunomodulatory treatments, or autoimmune/immunodeficiency diseases. Patient demographics and allergic indicators are summarized in Table S1. All donors were recruited with written consent and ethical approval from Hamilton Integrated Research Ethics Board (HiREB).

### Mice

Female C57BL/6 mice (5-8 weeks old) were purchased from Charles River Laboratories. 4get/KN2 mice were a generous gift from Dr. Irah King (McGill University). Mice were epicutaneously sensitized to- and challenged with crude peanut extract (CPE, Greer), as previously described (*14*). All experimentation was in compliance with McMaster University Research Ethics Board.

### Antibody Production and Administration

A hybridoma secreting monoclonal rat IgG2a anti-mouse IL-4Rα (M1) was generously provided by Dr. Fred Finkelman (Cincinnati Children’s Hospital, Cincinnati, OH). Hybridomas were expanded in tissue culture flasks with serum-containing media and then transferred to a spinner flask in serum-free media for up to 10 days. Antibodies were purified from culture supernatant with Protein G Sepharose (GE Healthcare). One mg anti-IL-4Rα or an isotype control (rat IgG2a anti-trinitrophenol; Bio X Cell) was injected intraperitoneally the day before subcutaneous re-exposure. IgG-mediated anaphylaxis was blocked with rat IgG2b anti-mouse CD16/32 (Bio X Cell), as previously published (*15*).

### PBMC Isolation and Storage

Up to 80 mL of peripheral blood was collected into heparinized tubes (BD). PBMCs were isolated via Ficoll-Paque (GE Healthcare) density gradient centrifugation, filtered, and resuspend in either enriched RPMI medium for culture, or in 10% dimethyl sulfoxide (DMSO)-supplemented fetal bovine serum (FBS) to be frozen at -80°C overnight in appropriate freezing containers and subsequently in liquid nitrogen for long-term storage. PBMCs used in this study were frozen for 0-6 months before usage.

### PBMC Culture

Before culture, PBMC viability was assessed using automated viable cell counting (ADAM MC, Digital Bio). Fresh PBMCs were >95% viability and freeze-thawed PBMCs were >90% viability. Cells were resuspended in enriched RPMI 1640 (Gibco) supplemented with 10% heat-inactivated human AB serum (Corning), 10 mM HEPES, 0.1 mM non-essential amino acids, 1 mM sodium pyruvate, 55 µM 2-mercaptoethanol, 1% L-glutamine, and 1% penicillin-streptomycin. Cells were plated in at least duplicate at a density of 1.5×10^6^ per mL in 24-well plates for up to 12 days. On days 4 and 8, 1 mL of cell-free supernatant was collected, stored at - 80°C, and replaced with 1 mL enriched medium. As a positive control for IgE production, cells were stimulated with 68.7 ng/mL (8000 IU) rhIL-4 (Sigma-Aldrich) and 5 µg/mL anti-CD40 (BioXCell) at the start of culture. As a negative control, cells were incubated in medium alone. Allergen-stimulated cells were cultured with 2.5 ng/mL CPE. Cells incubated with neutralizing antibodies against IL-4Rα and keyhole limpet antigen (KLH, IgG2a isotype control for IL-4Rα antibody) received 50 μg/mL in the presence or absence of CPE.

### ELISA and ELISpots

PN-specific mouse IgG1, IgE, and total human IgE ELISAs were performed as previously described (*3, 20*). For the detection of human PN-specific IgE antibodies, MaxiSorp plates (Thermo Scientific) were coated overnight at 4°C with 4 μg/mL CPE in carbonate-bicarbonate buffer. Wells were blocked with 5% skim milk powder in PBS for 2 hours followed by 3 washes with 0.05% Tween-20. Samples were plated and incubated overnight at 4°C. Supernatant samples were lyophilized (Modulyo Freeze Dryer, Thermo Scientific) overnight and were reconstituted (10x concentrated) in 5% skim milk. After washing, biotin-conjugated goat anti-human IgE, cross-adsorbed (polyclonal; Invitrogen) was added at 0.25 μg/mL. Following a 2 hour incubation, wells were washed and streptavidin-HRP (BD) was added for 1 hour. ELISAs were developed with 3,3’,5,5’-Tetramethylbenzidine (Sigma) and optical densities were acquired with a Multiskan™FC (Thermo Scientific).

Total IgE, PN-specific IgA, and PN-specific IgG ELISpots were performed with commercially available kits (Mabtech), as per manufacturer’s recommendations.

### Cytokine Analysis

Cytokines in PBMC culture supernatant were assayed using MILLIPLEX Immunology Multiplex Assays (HCYTOMAG-60K, Millipore Sigma) and analyzed using MAGPIX XMAP Technology (Luminex). Supernatants from day 4 of culture were assayed for either IL-4, IL-5 and IL-13 or a discovery panel including: sCD40L, EGF, Eotaxin/CCL11, FGF-2, Fit-3 ligand, Fractalkine, G-CSF, GM-CSF, GRO, IFN-alpha2, IFN-gamma, IL-1alpha, IL-1beta, IL-1Ralpha, IL-2, IL-3, IL-4, IL-5, IL-6, IL-7, IL-8, IL-9, IL-10, IL-12(p40), IL-12(p70), IL-13, IL-15, IL17A, IP-10, MCP-1, MCP-3, MDC (CCL22), MIP-1alpha, MIP-1beta, TGF-alpha, TNF-alpha, TNF-beta, VEGF. Cytokines detected above background (non-stimulated wells) and falling within the detectable range (3.2-10000 pg/mL) were included in statistical analyses.

### Flow Cytometry and Bulk Sorting

Cultured PBMCs or single cell suspensions from murine tissues were plated in PBS containing 0.5% Ethylenediaminetetraacetic acid (EDTA) and 0.5% bovine serum albumin (BSA) prior to extracellular staining. Cells were incubated with antibody cocktail and Fixable Viability Dye eFluor780 (eBioscience) for 30 minutes on ice. Fluorescently-conjugated antibodies used in flow cytometric analyses are listed in Table S4. For apoptosis analysis, cells were stained with Annexin V-PE in Annexin V binding buffer (BioLegend). Data were acquired with an LSRFortessa (BD). Bulk cell sorting was performed on a MoFlo XDP Cell Sorter (Beckman Coulter) prior to single-cell sorting. All data were analyzed with FlowJo v.10 software (TreeStar).

### Single-Cell RNA-Sequencing

Single Cell 3’ RNA Sequencing libraries were prepared using Chromium Single Cell V3 Reagent Kit and Controller (10X Genomics). Libraries were sequenced on HiSeq 4000 instruments (Illumina). The sequenced data were processed using Cell Ranger pipeline (v 3.0.1) by 10X Genomics (https://www.10xgenomics.com). Sequencing reads were aligned to the human transcriptome (GRCh38) followed by filtering and correction of cell barcodes and Unique Molecular Identifiers (UMIs). Reads associated with retained barcodes were quantified and used to build expression matrices.

The scRNA-seq R package Seurat (v3.1.1) was used for pre-processing, quality control and downstream analyses (*21*). Genes not detected in at least 3 cells were filtered out. Low quality cells expressing very few genes, and potential doublets or multiplets expressing aberrantly high number of genes (>5000) were excluded. Dying or stressed cells with a high percentage of transcripts (>20%) mapping to mitochondrial genes were removed. Samples from the four libraries were merged for subsequent clustering and visualization. Data were normalized using the log normalization method with a scaling factor of 10,000. Highly variable genes were selected using the mean-variance method and data were scaled to regress out the effect of nUMIs and percent mitochondrial genes. Dimensionality reduction was performed using principal component analysis (PCA) and the most significant clusters were chosen for subsequent clustering. Graph-based clustering was implemented by calculating k-nearest neighbors, followed by modularity optimization to clusters cells (FindClusters function). Non-linear dimensionality reduction and visualization was performed using UMAP (Uniform Manifold Approximation and Projection). Clusters were annotated on the basis of canonical markers and differential gene expression testing was used to determine a gene set signature for each cluster using the Wilcoxon Rank Sum test.

### Quantitative RT-PCRs

RNA was isolated from cultured PBMCs with RNeasy Mini Kit (QIAGEN) and was quantified with NanoVue Plus (GE Healthcare). cDNA was synthesized with Maxima First Strand cDNA Synthesis Kit (Thermo Scientific), as per manufacturer’s recommendations. Immunoglobulin epsilon (ε) heavy chain transcripts were amplified (StepOnePlus Real-Time PCR System, Applied Biosystems) using custom TaqMan gene expression kits (Applied Biosystems) with previously developed primer pairs and probes (*22*). Data was analyzed by ΔΔCT, to represent data as fold change relative to 18S housekeeping gene (Applied Biosystems) and unstimulated (media only) control.

### Statistical Analysis

Data were analyzed with Prism 8 (GraphPad Software). Comparisons were drawn using one- or two-way ANOVA, Student’s *t*-test, or Pearson r. Benjamini-Hochberg (BH) correction for multiple testing was used in analysis of cytokine discovery panel. Data were considered significant when *p* < 0.05.

## Acknowledgments

We kindly thank F. Finkelman and I. King for providing reagents and mice, respectively, T. Freitag and E. Avilla for help with patient recruitment, A. Dvorkin-Gheva and E. Al-Chami for assistance in statistical analyses. and M. Subapanditha for assistance with cell sorting.

## Author contributions

K.B. and P.S. designed and performed the experiments, analyzed data, and wrote the manuscript. S.V. performed the scRNA-seq analysis. O.B., S.M., S.G., M.D., A.F., T.W., J.W., and Y.E. assisted with experiments. D.C., R.J-S., S.E., and C.R. provided intellectual input and edited the manuscript. S.W. and M.J. raised funding. M.J. supervised the project and edited the manuscript.

## Data and materials availability

All data is available in the main text or the supplementary materials.

## Supplementary Materials

Figs. S1 to S9

Tables S1 and S4

