## Supplementary Materials for "Interrupting reactivation of immunological memory reprograms allergy and averts anaphylaxis" (from another version of this preprint: version 1, 2020-03-05)

**This PDF file includes:**

Figs. S1 to S9
Tables S1 and S4
Captions for Tables S2 and S3

**Other Supplementary Materials for this manuscript include the following:**

Tables S2 and S3 (Excel format)

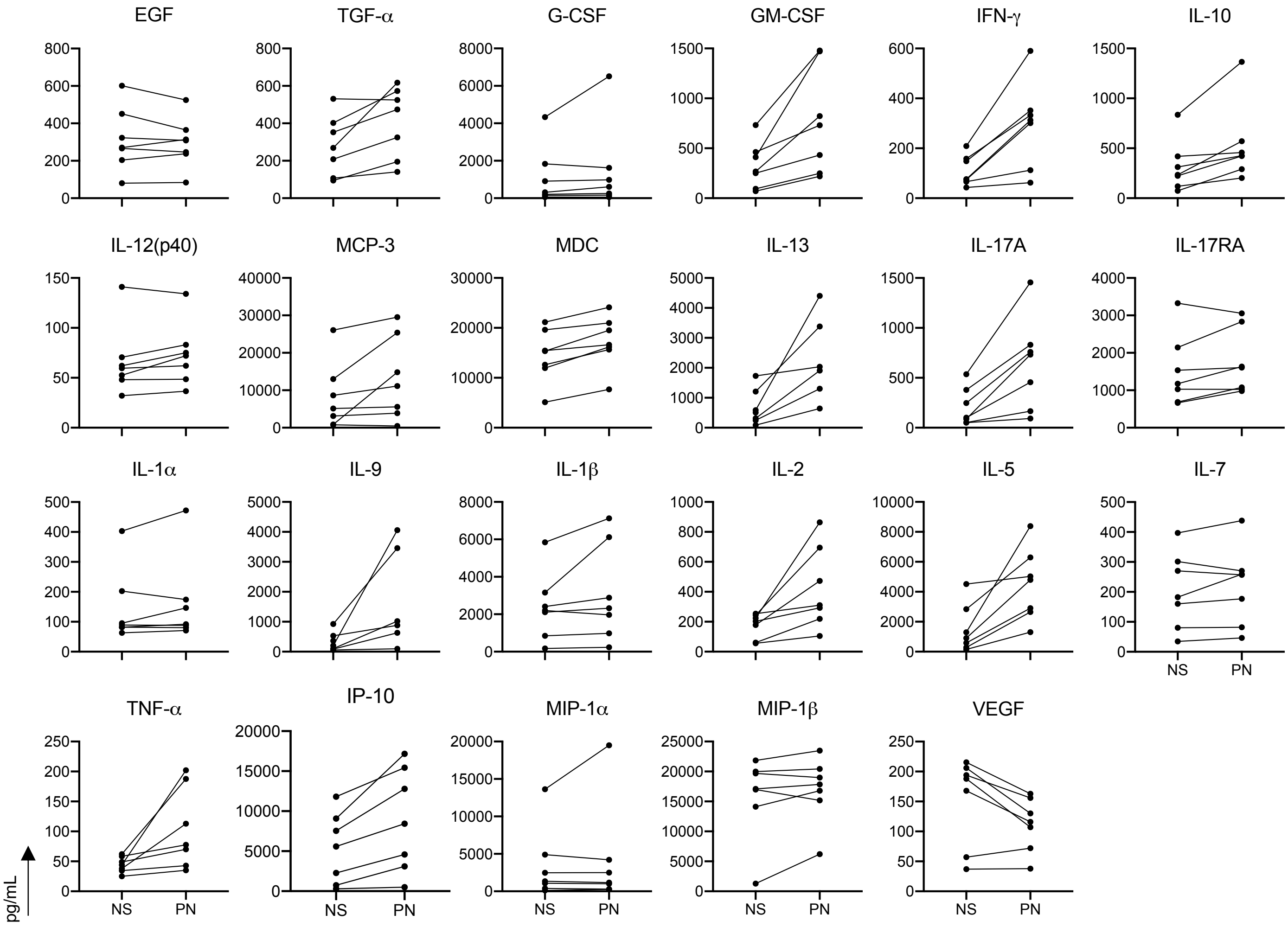


**Fig. S1.**

Cytokines detected (above background) with 38-analyte array in NS or PN-stimulated PBMCs from PN-allergic donors (n=7). Each line represents an individual donor.

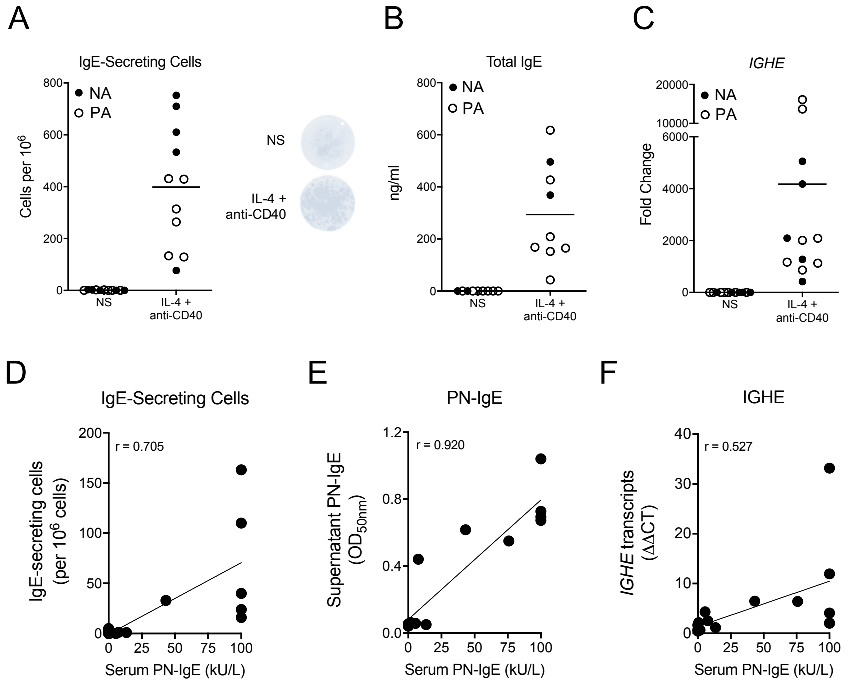


**Fig. S2.**

(A) Enumerated IgE-secreting cells detected at end of culture with representative images. (B) Total IgE measured from culture supernatant. (C) Fold change of *IGHE* mRNA expression relative to housekeeping gene (*18S*) and non-stimulated control. Data are representative of 2-5 non-allergic donors (NA; filled circle) and 6-7 PN-allergic donors (PA; open circle) and plotted as individual data points with mean (horizontal bar). (D-F) Relationship between serum PN-specific IgE levels (kU/L) and IgE readouts in 5 non-allergic donors and 7-11 PN-allergic donors. Linear relationship was calculated with Pearson’s r.

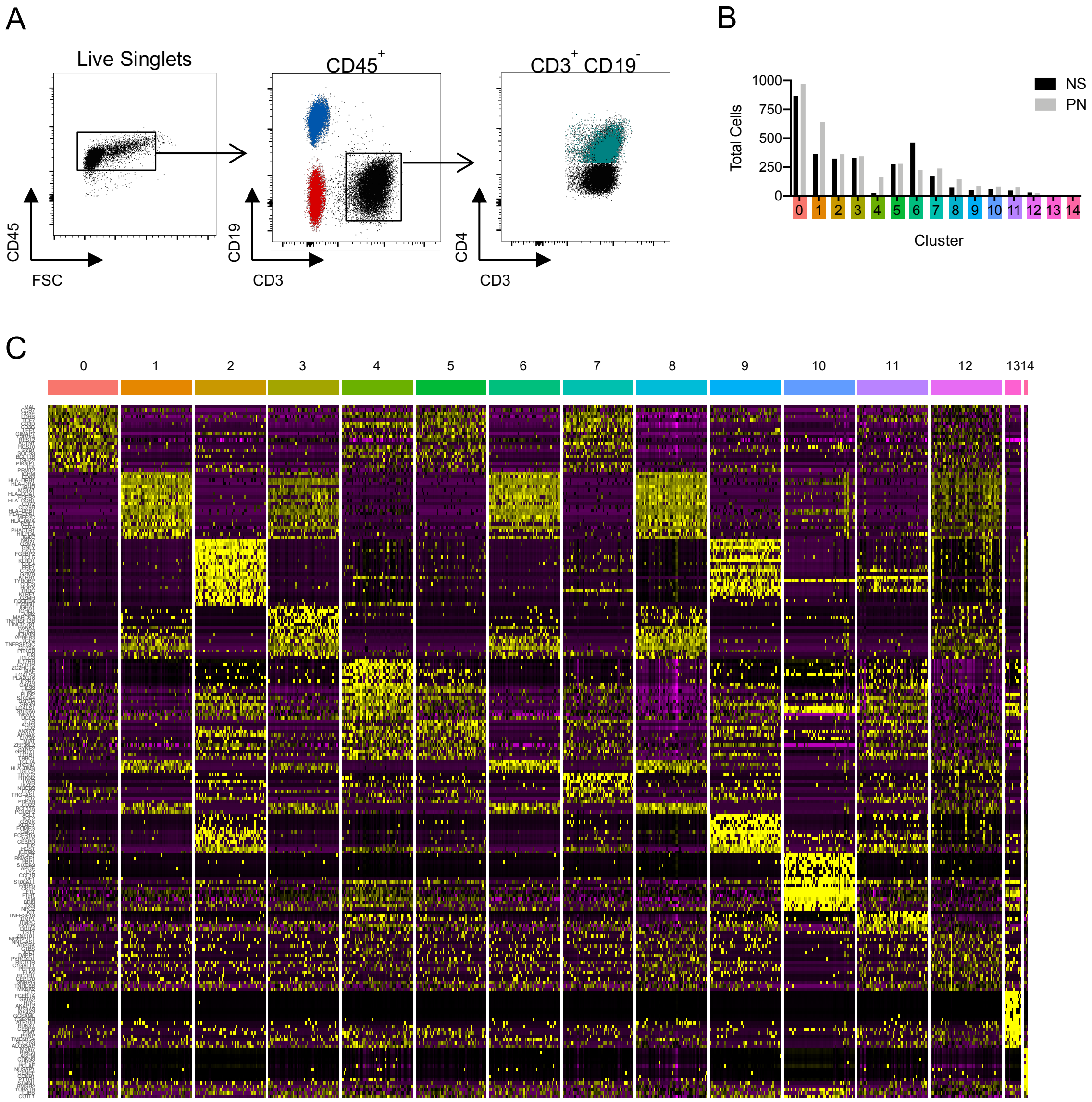


**Fig. S3.**

(A) Gating strategy for sorting of B cells, CD4$^+$ T cells, and non-T/B cells for single-cell RNA-sequencing. Sorted populations were pooled at the approximate frequencies: 40% CD4$^+$ T cells, 40% B cells, 20% non-T/B cells. (B) Total cells in each cluster of Fig. 1G. (C) Heatmap of top 20 differentially expressed genes in leukocyte clusters from Fig. 1G.

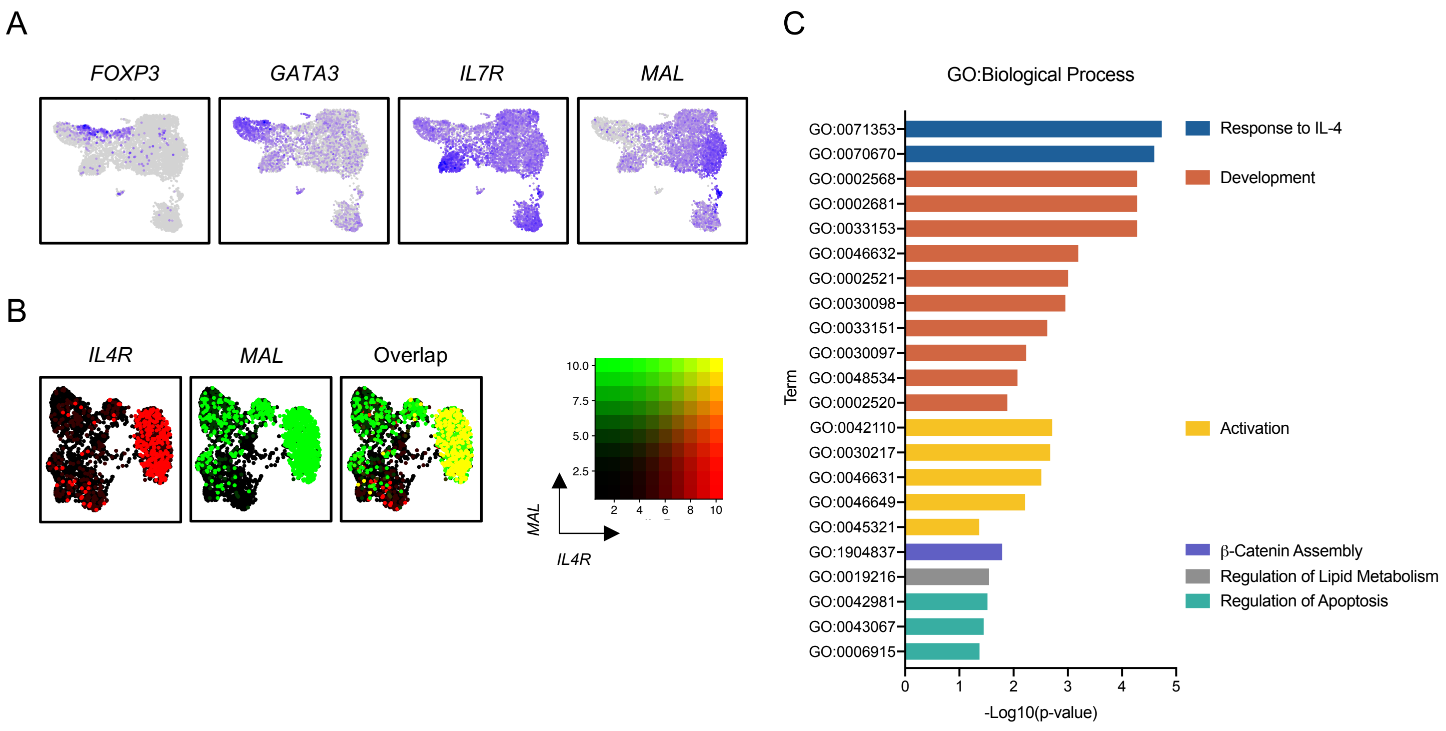


**Fig. S4.**

(A) UMAP visualization of T cell clusters with overlay of gene expression used for T cell subclustering in Fig. 1I. (B) Feature plots depicting *IL4R* and *MAL* expression and co-expression in T cell subclusters. (C) Pathway analysis (G:Profiler) of differentially expressed genes (excluding ribosomal genes) in $MAL^{hi}$ cluster.

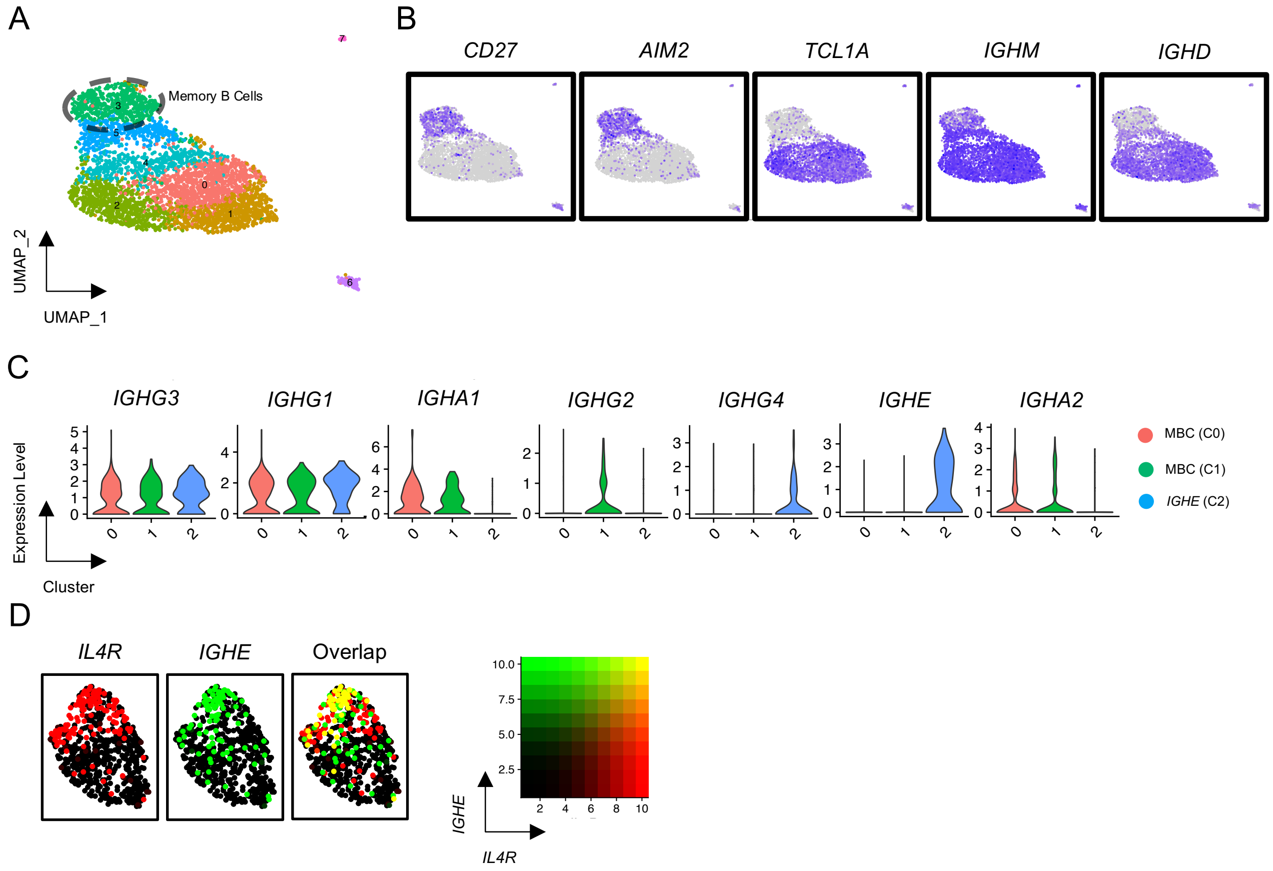


**Fig. S5.**

(A) UMAP of total B cells. (B) Feature plots of key genes used to subcluster MBCs in Fig. 1L. (C) Violin plots of IgH gene expression in MBC subclusters. (D) Feature plots depicting *IL4R* and *IGHE* expression and co-expression in MBC subclusters.

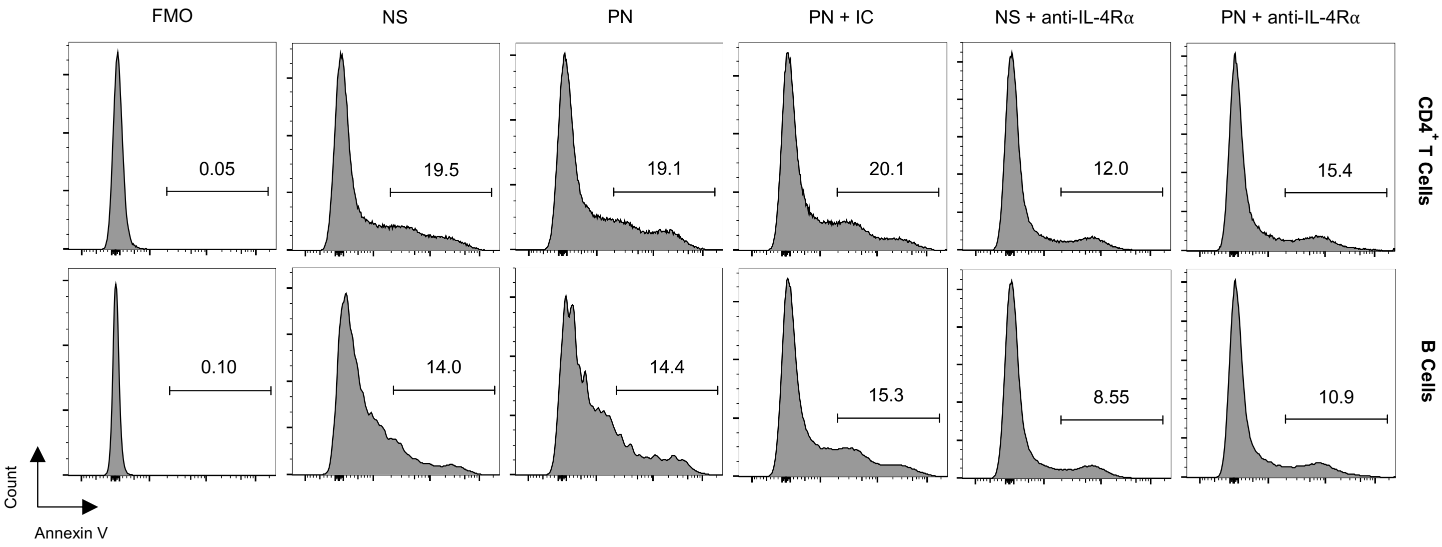


**Fig. S6.**

Representative histograms and mean frequency of Annexin $V^{+}$ $CD4^{+}$ T cells and B cells assessed by flow cytometry at end of culture.

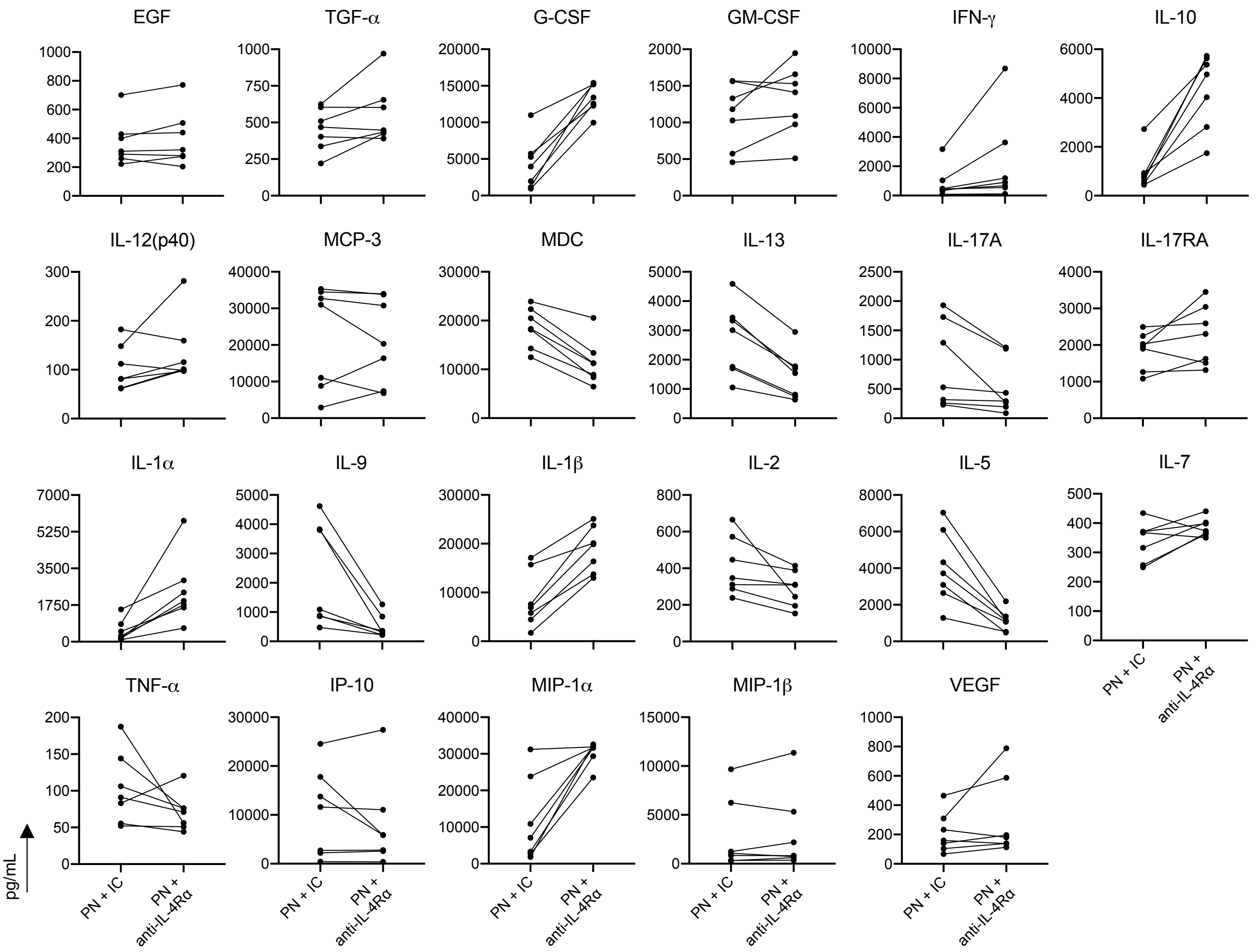


**Fig. S7.**

Cytokines detected (above background) with 38-analyte array in anti-IL-4Rα- or IC-treated PN-stimulated PBMCs from PN-allergic donors (n=7). Each line represents an individual donor.

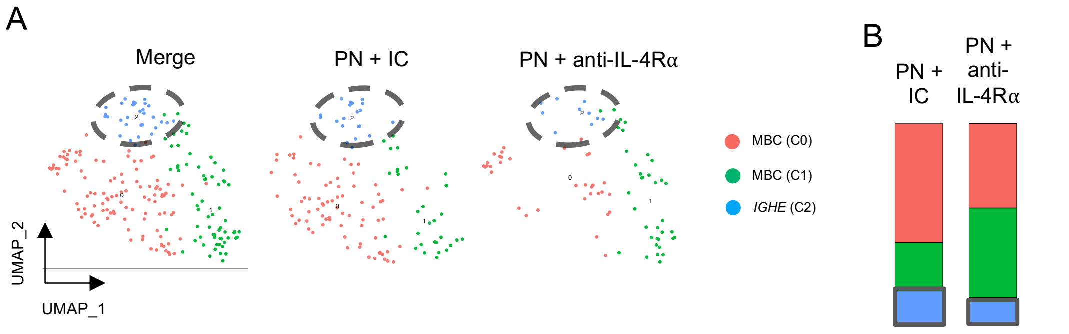


**Fig. S8.**
(A) UMAP of subclustered memory B cells. (B) Stacked bars represent subcluster frequency in A.

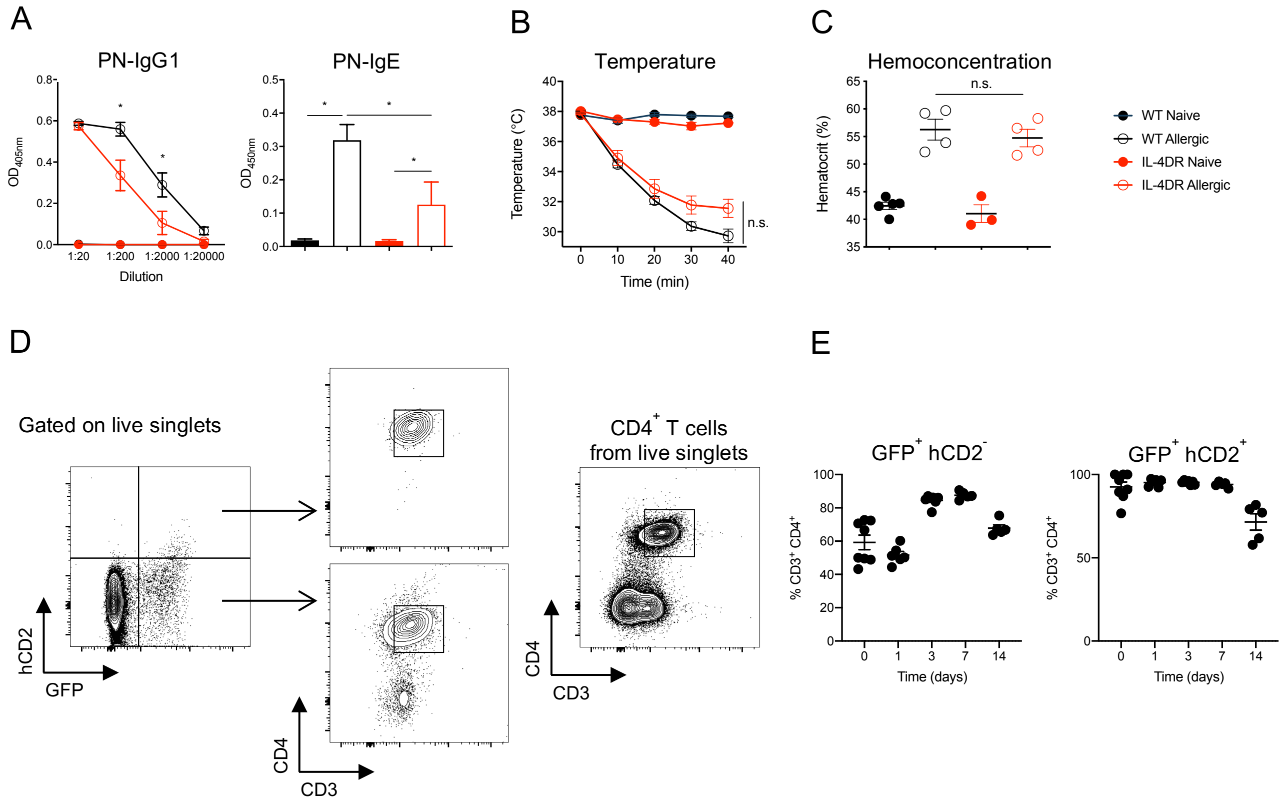


**Fig. S9.**

(A) Serum PN-specific IgG1 and IgE following sensitization of 4get/KN2 mice. (B) Core body temperature at 10-minute intervals during systemic challenge. (C) Hemoconcentration at 40-minutes post-challenge. (D) Representative contour plots of *IL4*-transcribing and IL-4 secreting cells and subgating on CD4$^{+}$ T cells. (E) Frequency of CD4+ T cells in *IL4*-transcribing and IL-4-secreting cells. Data are representative of 1-2 experiments with 3-5 mice per experimental group, plotted as mean ± SEM or median (I). **P*<0.05.

**Table S1.**
Patient demographics.

| Donor ID | Sex | Age | Serum PN-IgE (kU/L) | SPT (S-H-PN; mm) | Clinical history of PN allergy |
|---|---|---|---|---|---|
| P001 | M | 21 | <0.1 | 2-5-2 | No |
| P007 | M | 23 | <0.1 | 1-4-1 | No |
| P009 | F | 24 | <0.1 | 1-4-1 | No |
| P026 | M | 20 | <0.1 | 1-5-1 | No |
| P021 | M | 21 | <0.1 | 1-4-1 | No |
| P011 | M | 19 | 0.66 | 1-5-7 | Yes |
| P016 | M | 21 | 1.66 | 1-4-8 | Yes |
| P024 | M | 61 | 5.54 | 1-4-10 | Yes |
| P020 | M | 26 | 7.56 | 1-4-12 | Yes |
| P006 | F | 23 | 13.48 | 1-4-5 | Yes |
| P017 | M | 22 | 43.28 | 1-4-10 | Yes |
| P034 | M | 22 | 75.84 | 1-4-7 | Yes |
| P040 | F | 12 | >100 | 0-4-10 | Yes |
| P041 | F | 13 | >100 | 1-4-9 | Yes |
| P042 | F | 14 | >100 | 0-3-8 | Yes |
| P027 | M | 13 | >100 | 1-4-5 | Yes |
| P043 | M | 10 | >100 | 0-4-10 | Yes |

**Table S4.**
FACS antibodies.

| Target | Species Reactivity | Clone | Conjugate | Source |
|---|---|---|---|---|
| CD2 | Human | RPA-2.10 | APC | BD Biosciences |
| CD3 | Mouse | 17A2 | BV510 | BioLegend |
| | Human | HIT3a | FITC | BioLegend |
| CD4 | Mouse | RM4-4 | PerCP/Cy5.5 | BioLegend |
| | Human | OKT4 | BV510 | BioLegend |
| CD16/32 | Mouse | 93 | Unconjugated | BioLegend |
| CD19 | Mouse | 6D5 | PerCP/Cy5.5 | BioLegend |
| | Human | SJ25C1 | BV421 | BioLegend |
| CD44 | Mouse | IM7 | BV650 | BioLegend |
| CD45 | Human | HI30 | AF700 | BioLegend |
| CD62L | Mouse | MEL-14 | BV421 | BioLegend |
| CXCR5 | Mouse | L138D7 | PE, BV711 | BioLegend |
| PD-1 | Mouse | RMP1-30 | PE/Cy7 | BioLegend |
| Fixable viability dye | - | - | eFluor 780 | eBioscience |

**Table S2.**
List of differentially expressed genes in T cell subclusters. (Excel format)

**Table S3.**
List of differentially expressed genes in MBC subclusters. (Excel format)
